# Emergence of large-scale polar microtubule swarms for dense molecular transport

**DOI:** 10.64898/2026.07.06.736790

**Authors:** Meisam Zaferani, Ned S. Wingreen, Howard A. Stone, Sabine Petry

## Abstract

Microtubules (MTs) and their motor proteins collectively harness chemical energy to generate mechanical work, driving some of the most coordinated self-organized dynamics in living cells. The unique properties of these molecules also make them versatile building blocks of cytoskeletal active matter and biomimetic nanomachines that recapitulate cellular motility, emergent pattern formation, and motor-driven transport. However, these canonical systems use MTs of fixed length and do not incorporate the natural ability of MTs to grow and regenerate. Here, we go beyond these limits by using dynamic self-amplifying branched MT networks. Driven by kinesin-1 and cytoplasmic dynein activity, surface-gliding branched MT bundles undergo swarming that yields large-scale collective MT architectures with several sought-after features. They are polar and orientationally aligned, dense, span millimeter scales, and persist over hours. We then show that these features enable molecular transport along the swarm at unprecedented capacities, with up to six million motor complexes walking in parallel across millimeter-scale distances over hours. Our results introduce a new regime in cytoskeletal active matter in which the interplay between motor-driven activity and filament generation via branching leads to emergent polar order in proliferating swarms. Such emergent polarity makes these swarms suitable for engineering scalable transport nanotechnologies and programmable soft materials.

## 1. Introduction

Active matter is composed of energy-consuming building blocks that generate local motion and mechanical stresses, leading to far-from-equilibrium collective behaviors^1^. Examples of active matter at the microscale include self-propelled colloidal particles^2–4^, assemblies of microswimmers^5–11^, cytoskeletal architectures^12–14^, and collections of cells that form tissues^15^. The emergent collective behaviors in these systems, such as hierarchical self-organization^16^, dynamic pattern formation^17^, and spontaneous motion^18,19^, have attracted considerable attention from a nonequilibrium physics perspective^20–23^. Moreover, harnessing these emergent phenomena enables bio-inspired materials and technologies^24–26^.

Cytoskeletal filaments are central to biological active matter. Actin filaments and microtubules (MTs) provide structural scaffolds, while molecular motors generate directed forces that drive motion along the filaments. *In vitro* reconstituted systems of actin–myosin^27,28^ and MT–motor^29–31,16^ have therefore emerged as powerful tools for probing emergent phenomena in active matter and identifying the physical principles underlying nonequilibrium self-organization. In particular, the collective interactions of MTs with kinesin and dynein motors have also enabled the engineering of programmable materials^32–37^, molecular transport nanotechnologies^38–45^, as well as nanorobotics^46–49^.

Most existing approaches rely on using stabilized MTs as fixed-length tracks for motors, while MT filament length and population remain static^50^. By contrast, MTs in cells are not only dynamic, but are also continuously regenerated in an organized manner, leading to continuous remodeling of the MT architecture in cells. As a result, current MT-based active matter systems do not harness some key nonequilibrium features of the cell’s cytoskeletal toolkit.

With this motivation, we build upon the cell’s endogenous MT machinery to explore a new regime in cytoskeletal active matter. We focus on the ability of MTs to self-amplify through branching MT nucleation, in which new filaments are generated from the sides of existing MTs^51,52^. Branching MT nucleation has been observed during cell division^53^ and in developing neurons^54^, and its potential for use in nanotechnology^55,56^ has been previously demonstrated. Here, integrating branching MT nucleation with molecular motor activity, we construct a new cytoskeletal active matter system. The collective properties of this system emerge from swarming dynamics among MT bundles and have no precedent in canonical active matter.

## 2. Results

### 2.1. Swarming among branched MT bundles

To access the cell’s own MT machinery, we used *Xenopus laevis* egg extract, which provides all the raw material in a cell-free environment to form the MT cytoskeleton, while being accessible for biophysical and biochemical manipulations (Fig. 1a). Extract was supplemented with fluorescently labeled tubulin and GFP-tagged end-binding protein 1 (EB1) to visualize MTs and their growing plus ends, respectively. Upon introducing the supplemented extract in a flow cell and observing it via total internal reflection (TIRF) microscopy, individual MTs nucleated on the glass coverslip with density of 20–40 MTs per 200 × 200 µm² over 30 min (*N* = 5) and continued to grow with speed of *v =* 7 ± 3 µm/min (*N* = 15).

**Figure 1.**
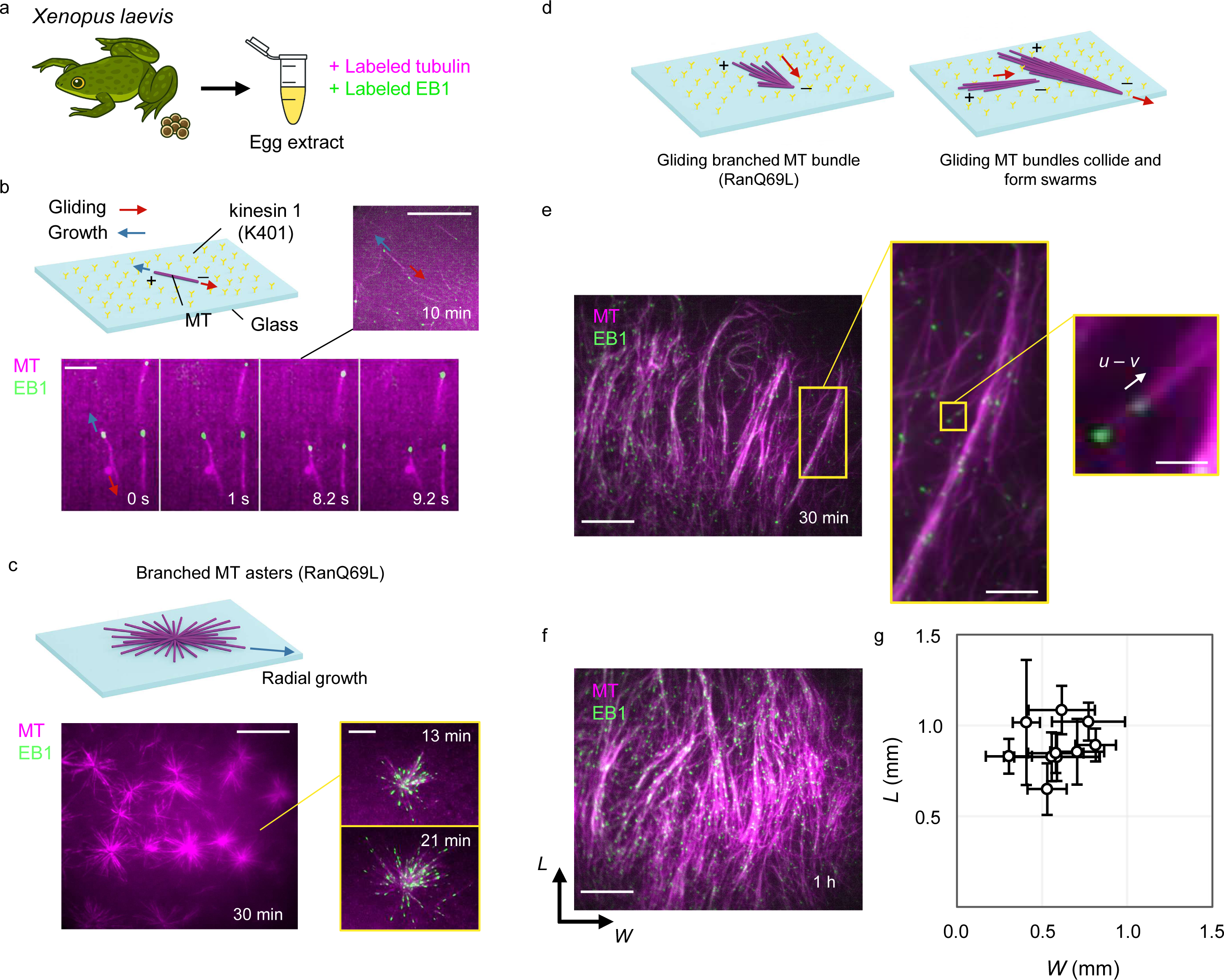
Formation of polar MT swarms. a) Xenopus egg extract supplemented with fluorescently labeled tubulin and GFP-tagged EB1 to visualize MTs and their growing plus ends. b) In the absence of RanQ69L, individual MTs nucleate on the coverslip and grow while gliding on K401-coated surfaces. c) RanQ69L induces stationary branched MT asters on non-motor-coated surfaces. EB1 comets in the right panel at two time points show radial growth of the aster. d) On K401-coated surfaces, the radial symmetry of branched MT asters is broken, leading to the formation of gliding MT bundles that collide and self-organize into MT swarms. e) An aligned MT network forms approximately 30 min after reaction initiation. EB1 comets reveal uniform MT polarity within aligned bundles, and the apparent EB1 speed (*u − v*) is used to infer gliding contribution. f) Dense polar MT swarms form within 1 h of reaction initiation. g) Swarm size quantified by length (*L*) and width (*W*) (*N* = 10 from 5 independent replicates). Data are shown as mean ± SD. Scale bars: 30 µm in b (top), 10 µm in b (bottom), 50 µm in c (left), 20 µm in c (right), and 50 µm in e and f.

We first asked how motor-driven gliding by kinesin, which allows local interactions (collisions) among these individual dynamic MTs, affects their self-organization. To do so, we immobilized purified truncated kinesin-1 (K401) on the coverslip and added the supplemented extract. As a result, MTs glide on the surface toward their minus ends, while simultaneously growing from their plus ends (Fig. 1b, Movie S1). Measurements of their growth (*v =* 8 ± 2 µm/min) and gliding speeds (*u* = 28 ± 4 µm/min) indicate that gliding is substantially faster than growth (Fig. S1a). We observed that collisions rarely lead to alignment but instead result in crossing events, with no emerging swarming behavior (Fig. S1b, Movie S1). This is consistent with previous studies using stabilized MTs in a purified system^31^ and indicates that MT growth per se does not lead to swarming.

A more important ability of MTs in *Xenopus* egg extract is to regenerate themselves via branching MT nucleation, in which new MTs nucleate at shallow angles on existing MTs, preserving polarity and enabling exponential amplification^57^. We induced branching MT nucleation by adding constitutively active Ran (RanQ69L), which releases branching factors that, together with MT nucleators, bind existing MTs, initiate new branch sites, and promote MT branching. If motor activity is inhibited, addition of RanQ69L leads to the formation of asymmetric branched MT networks (Fig. S2a), in which exponential and polarized MT amplification are clearly visible, as reported previously^51^. However, when motors are active in the extract, wild-type dynein, together with dynactin and NuMA^58^, focus the minus ends of MT branches to form radially branched MT asters (Fig. 1c, Movie S2). These asters grow outward but do not translate on the surface.

To investigate how motor-driven surface gliding affects these branched MT asters, we next coated the coverslip with K401 before addition of the extract. Under this condition, surface gliding induces symmetry breaking of radial asters, converting them into gliding MT bundles (Fig. 1d, Movie S3). Based on EB1 localization at growing plus ends, we observed that surface gliding occurred toward the minus end of the bundle, while MTs within the bundles continued to grow in the plus-end direction and regenerate through branching MT nucleation. Unlike individual MTs, these gliding bundles physically interacted upon collision and exhibited collective surface gliding. Subsequently, we observed the emergence of large-scale surface gliding MT swarms with strong orientational order, in which inter-filament angles remained below 5° (Fig. 1e, Fig. S2b). The initial orientation of the swarms varied randomly across replicates, consistent with their emergence through symmetry breaking and collective dynamics. However, once established during the early stages (5–10 min), the swarm orientation remained largely stable over time, exhibiting only small fluctuations.

To quantify the generic features of a swarm, we first determined the polarity of its MTs and their growth speed by tracking the EB1 signal on growing MT plus ends. MTs were not only orientationally ordered but also shared common polarity (Fig. 1e, Movie S4). The relative plus-end speed (*u-v*), which corresponds to the gliding speed of EB1 comets in the swarm, was 16 ± 3 µm/min at 30 min after reaction initiation (Fig. S3). This measurement shows no significant difference from individual MTs (Fig. S1a), indicating that MT growth and gliding speeds are unchanged in the swarm. However, within approximately one hour, the gliding speed decreased to below 2 µm/min and the growth speed to below 3 µm/min, resulting in almost static architectures. By one hour, branching MT nucleation allowed these swarms to reach an average transverse linear density of 3–9 MTs per µm and an average surface density of (2.5 ± 0.5) × 10^3^ MTs per 200 × 200 µm² (Fig. 1f). To precisely quantify the sizes of these architectures, we fixed them with 0.25% glutaraldehyde and measured their length along the main axis of alignment (*L*) and width in the perpendicular direction (*W*), which indicated an immense size of ∼1 x 0.5 mm^2^ (Fig. 1g, Fig. S4).

These swarms combine several sought-after properties. They grow to millimeter-scale from nanometer-scale building blocks, exceeding their molecular components by a factor of 10^5^-10^6^. They share a common polarity across all MTs and maintain order for hours, far longer than most reconstituted active systems. Together with their high MT density, these properties define a distinct regime of cytoskeletal active matter with the potential for nanotechnology applications.

### 2.2 Dynamics of polar MT swarms

Next, we investigated the dynamics of these swarms. They propagate with a filamentous leading front that displays continuous gliding, followed by a dense interior region (Fig. 2a and b, Movie S5). To quantify the collective motion in the swarm, we used optical flow analysis to extract the local gliding directions both in the filamentous front (Fig. 2c, Movie S6) and in the dense interior region (Fig. 2d, Movie S6). Based on this analysis, we then computed the gliding direction of the swarm (Fig. 2e) at two time points (15 and 30 min). Additionally, we quantified the degree of order in the collective motion by computing the polar order parameter, *S* = ⟨cos(*θ* − ⟨*θ*⟩)⟩, where *θ* is the gliding angle (relative to the horizontal), and 〈·〉 denotes an average across the swarm. The order parameter is surprisingly high even at early stages (*S* ≈ 0.7 at 15 min) and increases to *S* ≈ 0.8 at 30 min as the swarm proliferates, indicating a unique form of polar self-organization (Fig. 2f). The nearly constant gliding direction, together with the high polar order, indicates coherence within the swarm.

**Figure 2.**
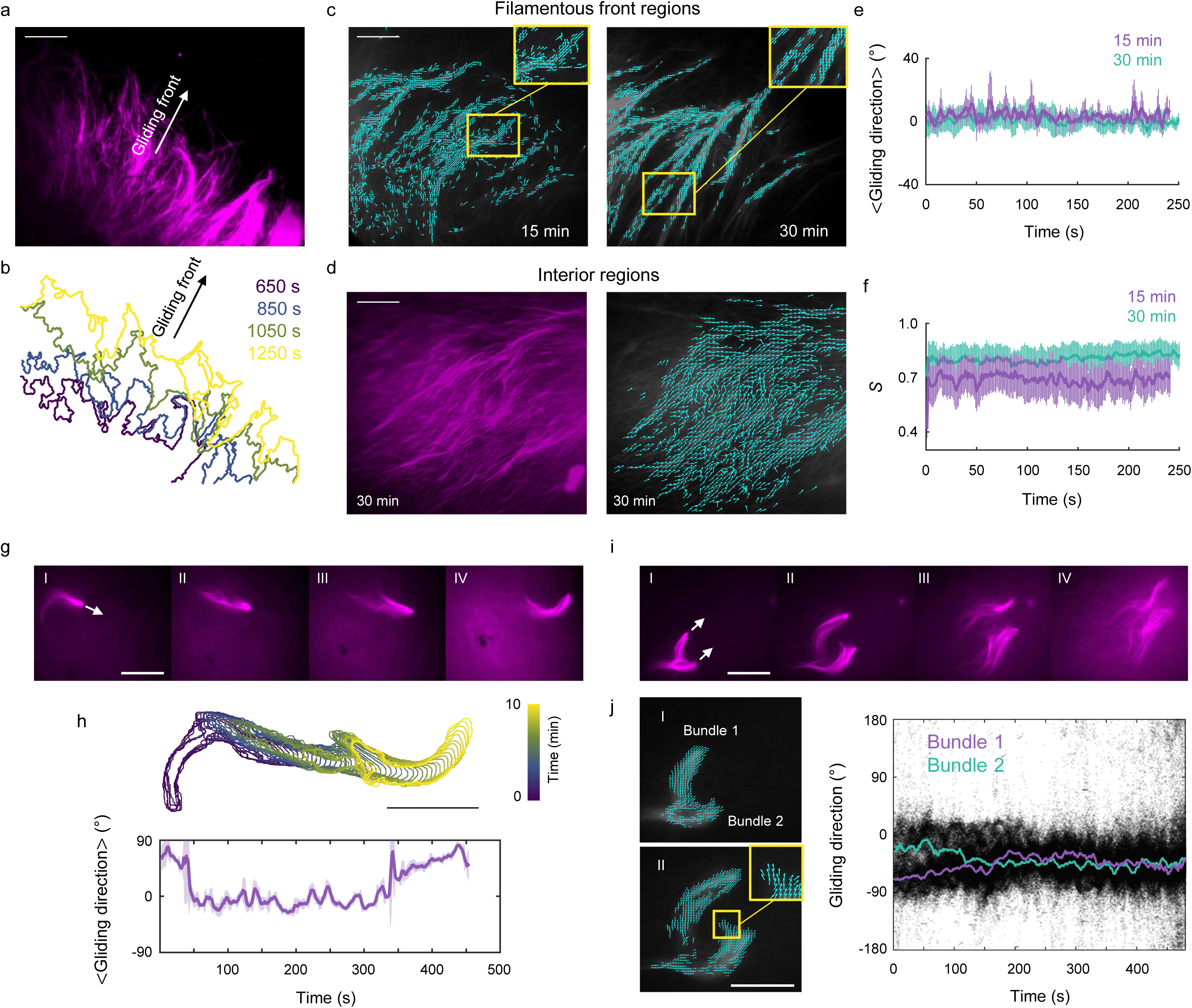
Dynamics of polar MT swarms. a) Filamentous morphology at the gliding front of the swarm. b) Overlay of gliding fronts color-coded by elapsed time. Scale as in a. c) Local MT gliding directions (from imaging at 1 frame per second) measured at the filamentous front, and d) within the interior region. e) Global swarm orientation measured 15 and 30 min after reaction initiation. f) Polar order parameter (*S*) quantified 15 and 30 min after reaction initiation. g) Representative images of a MT bundle remaining intact during gliding; white arrow indicates gliding direction. The interval between images is 90 s. h) Top: trajectory of the bundle shown in g, obtained by overlaying segmented bundle boundaries color-coded by elapsed time; persistence length is approximately 80 µm. Bottom: overall gliding direction of the bundle over time. i) Occasional division of gliding MT bundles followed by realignment; white arrows indicate gliding directions. The interval between images is 90 s. j) Left: gliding directions of daughter bundles after division (corresponding to I and II in panel i), demonstrating intrabundle coupling that maintains alignment. Right: quantification of gliding directions during and after bundle division; black dots indicate the gliding direction associated with each arrow, and lines represent the mean gliding direction for each bundle. Data are shown as mean ± SD where applicable. Scale bars: 50 µm for all panels except j (20 µm).

Alongside the formation of large-scale swarms, we also observed individual branched MT bundles gliding outside these swarms before merging into them. We analyzed these bundles as building blocks of the swarm, since understanding their gliding properties helps explain the coherence in the proliferating swarm. We noticed that individual bundles remained mostly intact while gliding (Fig. 2g, Movie S7) and moved persistently over distances of 60-80 µm before significantly reorienting (Fig. 2h). When reorientation occurred, the angular change was 50° ± 20° (*N* = 19). Additionally, we observed that gliding bundles occasionally divided (*N* = 2 out of 19). Surprisingly, despite splitting at an angle of ∼90°, the divided bundles subsequently glided parallel to one another, sometimes displaying intrabundle coupling during parallel gliding (Fig. 2i and j, Movie S7).

These observations show that individual bundles glide persistently over long distances and do not change direction by more than 90°. This persistence at the intrabundle level, together with aligning interactions among bundles, may further reinforce swarm coherence and order. Additionally, intrabundle coupling may enhance swarm robustness, which is crucial for coherency. Since such behaviors were not observed for individual MTs, we next investigated the underlying mechanisms responsible for this effect.

### 2.2. Branching MT nucleation and dynein activity are essential for swarm formation

Physical aligning interactions are a hallmark of emergent order in active matter. While individual gliding MTs did not display aligning interactions, gliding branched MT bundles aligned upon collisions and merged into larger composite and polar bundles with their leading (minus) ends pointed in the same direction (Fig. 3a and c, Fig. S5, Movie S8). Additionally, individual MTs were incorporated into the bundles upon collisions (Fig. 3b and c, Movie S8). These observations suggest that branching MT nucleation and dynein activity are necessary for alignment during collisions, a process that underlies the formation of polar MT swarms. We next tested this hypothesis.

**Figure 3.**
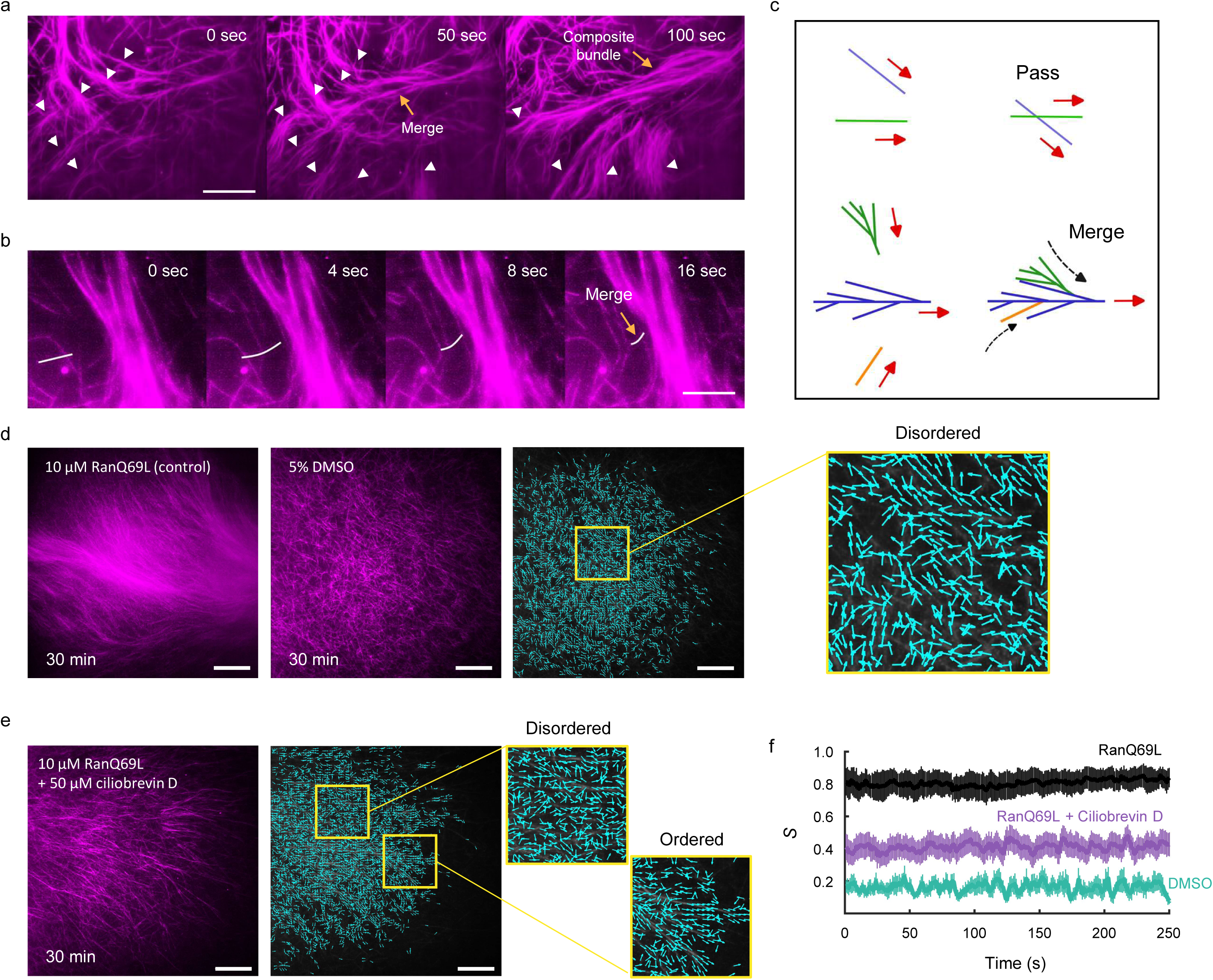
Branching nucleation and dynein activity drive swarm formation. a) Branched MT bundles align and merge upon collision, forming a composite bundle; white arrows indicate distinct branched MT bundles. b) Incorporation of an individual MT into a branched bundle. c) Schematic of MT–MT interactions: individual gliding MTs pass over one another (top), whereas individual MTs and branched bundles align and merge into larger branched composite bundles upon collision (bottom). d) Left: MT swarms formed in the presence of RanQ69L. Middle: dynamic individual MTs formed with 5% DMSO, which lacks branching MT nucleation. Right: local MT gliding directions corresponding to the middle panel, showing disordered gliding under 5% DMSO conditions. e) Left: dynein inhibition disrupts swarm formation. Right: MT gliding directions corresponding to the left panel, showing coexistence of ordered and disordered regions. f) Polar order parameter measured for MT swarms formed with RanQ69L, RanQ69L with dynein inhibition, and 5% DMSO. Data are shown as mean ± SD. Scale bars: 20 µm in a, 10 µm in b, 50 µm in d and e.

To assess whether branching MT nucleation is necessary for the emergence of these swarms, we eliminated branching MT nucleation by replacing RanQ69L with dimethyl sulfoxide (DMSO). Addition of 5% DMSO promotes spontaneous nucleation of individual MTs and their growth, yielding high densities of (2-4) × 10^3^ MTs per 200 × 200 µm² over 30 min (*N* = 3), but not in the branched form. Yet analogous to RanQ69L asters, these individual MTs formed asters on non-K401-coated surfaces (Fig. S6A), in agreement with previous reports^59^. On K401-coated surfaces, these individual MTs also glided on the surface, but did not exhibit aligning interactions upon collision (Fig. S6B, Fig. S7). Therefore, no MT swarming was observed, and gliding directions remained disordered (isotropic), as quantified by optical flow analysis (Fig. 3d). Consequently, the order parameter remained below 0.2, even at high MT densities (Fig. 3f, Movie S9). Thus, branching MT nucleation is essential for the emergence of MT swarms.

To assess the role of cytoplasmic dynein, which is known to promote pole focusing in MT assemblies, we induced branching MT nucleation using RanQ69L in the extract and on K401-coated surfaces. However, this was performed in the presence of 50 μM ciliobrevin D, a specific inhibitor of cytoplasmic dynein, to selectively suppress dynein activity^60^. We did not observe MT swarms with long-range order, but instead a mixture of regions exhibiting ordered (polar) and disordered (isotropic) MT gliding, as quantified by optical flow analysis (Fig. 3e, Movie S9). Consequently, the order parameter was ∼ 0.4 (Fig. 3f). These results suggest that while branching MT nucleation generates locally aligned MTs, dynein activity is also important for polar alignment among bundles and, consequently, for establishing long-range order in the swarm.

### 2.3. Molecular transport via swarms

In previously developed MT-based transport nanotechnologies, motor proteins were immobilized on surfaces and individual MTs acted as shuttles. Aligning the gliding direction of MTs in these approaches is, however, challenging, and efforts have been devoted to addressing this issue using external forces^61^. In contrast, our MT swarms exhibit emergent polar order, eliminating the need for external control. This emergent order, together with the dense and coherent organization demonstrated in previous sections, provides an intrinsic alignment mechanism that can be harnessed for enhanced transport efficiency.

Another mode of molecular transport involves motors carrying cargo as they move along MTs. However, transport on individual MTs, which are typically only several microns long, is insufficient for effective transport nanotechnologies^62^. We hypothesized that this limitation can be overcome in our millimeter-scale MT swarms, which in principle are capable of scaffolding large numbers of motors moving uniformly along densely packed, polar-ordered MTs within the swarm.

To test this idea, we assembled a standard fluorescent motor complex by incubating biotinylated K401 motors with streptavidin-conjugated quantum-dot nanoparticles (Fig. 4a). Upon addition of these complexes to the extract, 70–90% of them were active and distributed throughout the swarm (Fig. 4b, Movie S10). By tracking individual motor complexes and extracting kymographs of their motion, we found that over 90% of them indeed moved unidirectionally throughout the swarm, toward the plus end, with an average speed of 18 ± 4 µm/min (Fig. 4c and d, Fig. S8).

**Figure 4.**
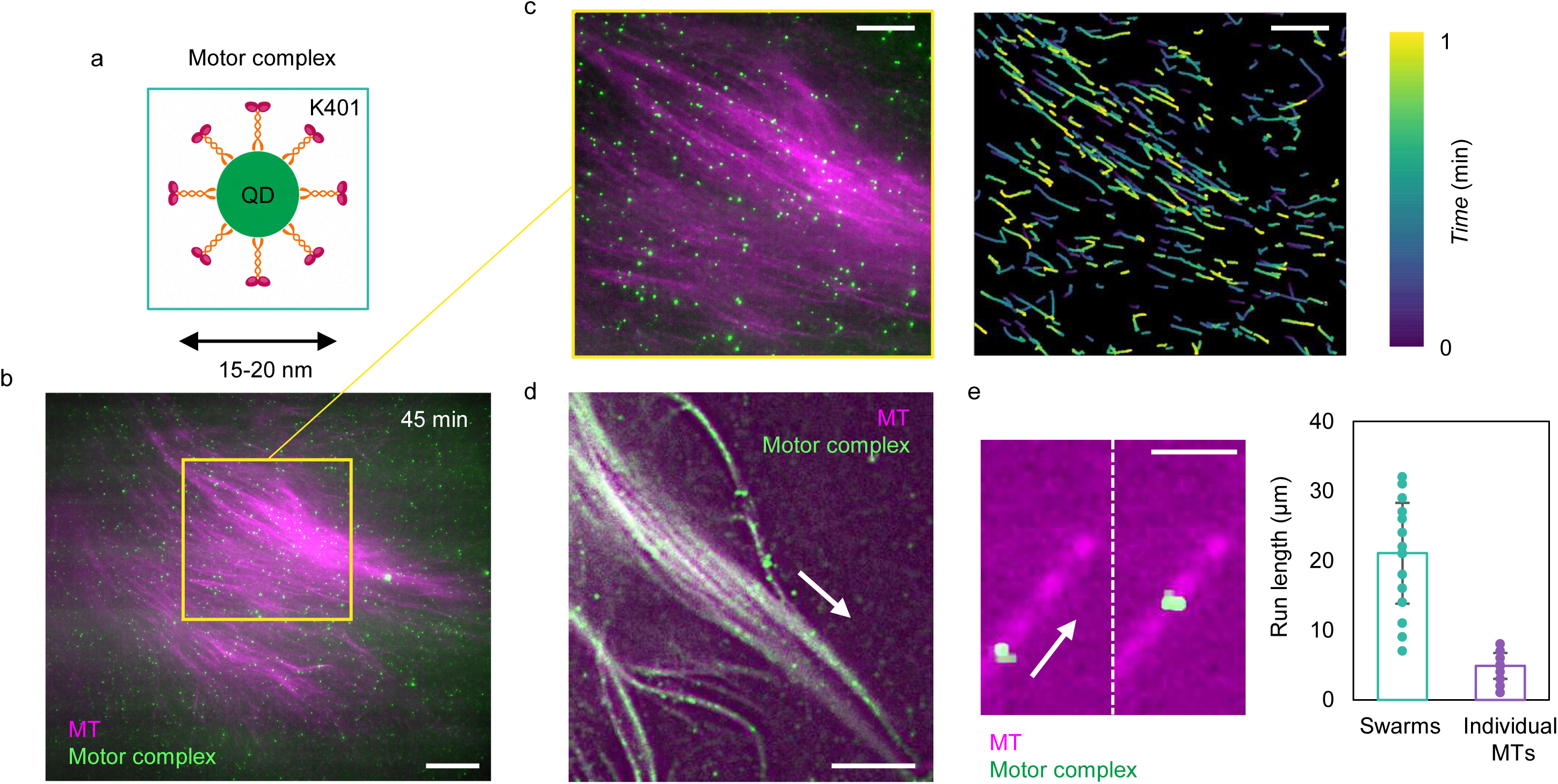
Dense transport of motor complexes via polar MT swarms. a) Motor complexes formed by conjugating streptavidin coated quantum dots to biotinylated K401 motors. b) Motor complexes move along polar swarms at densities of 3–12 complexes per µm², with 70–90% active motors (*N* = 10 independent replicates). c) Left: magnified region from b. Right: corresponding trajectories of motor complexes within the magnified region, color-coded by elapsed time; for better visualization of transport, only a subset (∼50%) of trajectories are shown. d) Transport of motor complexes at the rear of the swarm and toward the plus end. e) Left: transport of a motor complex on an individual MT. Right: minimum run length of motor complexes within swarms compared with run length on individual MTs (*N* = 30 motors from 3 independent replicates; *p* < 10⁻⁵). Scale bars: 50 µm in a, 25 µm in b and c, 10 µm in d, and 5 µm in e.

Molecular transport along MT swarms offers several advantages over transport on individual MTs. The density of motor complexes along the swarms (3-12 µm^-2^) was dramatically higher than that observed on individual MTs (∼10_-2_ µm^-2^), leading to the parallel transport of up to six million motor complexes within a single swarm. Additionally, we found that the minimum run length of motor complexes on the swarms, defined as the distance traveled before detachment, was at least fivefold greater than that on individual MTs (Fig. 4e, Fig. S9, Movie S11). These reported values represent lower bounds, because the run length on MT swarms exceeded the capacity to image longer movies, whereas this limitation did not apply to the measurements on individual MTs. These results demonstrate the unprecedented capacity of MT swarms for dense, large-scale, and directed molecular transport.

## 3. Discussion

Our results expand the scope of active matter by introducing a new type of emergent collective behavior. By combining motor activity in the bulk and on the surface, with filament generation through branching MT nucleation, we created dense, large-scale, and coherent polar swarms. Earlier MT-based active matter studies identified extensile active nematics^16^, contractile networks^32,63^, and vortex lattices^31,64^, and our results add coherent polar swarms as a complementary class. This work motivates future efforts to further explore the design space of cytoskeletal machinery in cells, aiming to uncover new classes of active matter and strengthen the links between active matter physics and biomolecular self-organization.

We found that aligning interactions, essential for swarm emergence, occurred among bundles and between bundles and individual MTs, but did not occur among individual MTs. This observation likely has several complementary explanations. Bundles have larger effective interaction interfaces, which enhance both steric effects and motor-mediated polar alignment between them. In addition, their increased rigidity and stronger collective adhesion to the surface increase the likelihood of physical interactions compared to individual MTs. Some MT-associated proteins released upon the addition of RanQ69L are known to bundle MTs and may therefore further enhance alignment of MTs. Moreover, biomolecular condensates have been shown to generate capillary forces between MTs^65^. Therefore, it is possible that TPX2, a key condensate involved in branching, also contributes to alignment. Finally, the larger size of bundles relative to individual MTs makes it plausible that hydrodynamic interactions contribute to their alignment as well.

This work also introduces polar MT swarms suitable for molecular transport nanotechnologies, where the emergent polarity and size overcome the key limitations of existing motor-based systems that depend on controlling the gliding directions of stable MTs^61,62^. Future efforts can build upon this system by optimizing transport efficiency and using diverse motor complexes to achieve different forms of transport. Additionally, leveraging micro- and nanofabrication^41^ techniques together with optical tools^32,38,49^ will enable programmable control of molecular transport. The dynamics of these swarms can also be harnessed to engineer soft materials and complex fluids for targeted applications. Designed to respond to external stimuli, such biomimetic materials would reconfigure autonomously and perform programmable mechanical tasks with molecular-scale precision.

## 4. Methods

### 4.1. Xenopus egg extract and protein purification

#### Xenhopus egg extract preparation

Eggs were obtained from adult *Xenopus laevis* females hormonally primed with human chorionic gonadotropin (hCG) to induce ovulation. Mature eggs were collected, dejellied, and cytosol was prepared from eggs naturally arrested in meiosis II, following modified previously described protocols^51^. Extracts were supplemented with fluorescently labeled tubulin (1 μM) to visualize MTs, GFP-EB1 (100 nM) to mark growing plus ends, and RanQ69L (10 μM) to induce branching nucleation. A 1% (v/v) energy mix (0.75 ml 1 M creatine phosphate, 1 ml 0.1 M ATP, 0.1 ml 1 M MgCl₂, 3.15 ml H₂O) was included to maintain ATP levels. Reactions were assembled on ice and maintained at 0–4 °C until injection into flow chambers for imaging.

#### Protein purification

EB1-GFP was expressed in *Escherichia coli* (Rosetta 2 DE3) at 37 °C for 4 h and lysed in 50 mM NaPO₄, pH 7.4, 500 mM NaCl, 20 mM imidazole, 2.5 mM PMSF, 6 mM β-mercaptoethanol (BME), 1 cOmplete EDTA-free protease inhibitor (Sigma), and 1,000 U DNase I (Sigma). Lysates were clarified and applied to a HisTrap HP 5 ml column (GE Healthcare), washed with binding buffer (50 mM NaPO₄, pH 7.4, 500 mM NaCl, 20 mM imidazole, 2.5 mM PMSF, 6 mM BME), and eluted with 50 mM NaPO₄, pH 7.4, 500 mM NaCl, 500 mM imidazole, 2.5 mM PMSF, 6 mM BME. Pooled fractions were further purified by gel filtration on a Superdex 200 pg 16/600 column in CSF-XB (10 mM HEPES, pH 7.7, 1 mM MgCl₂, 100 mM KCl, 5 mM EGTA) containing 10% (w/v) sucrose.

RanQ69L, bearing an N-terminal blue fluorescent protein (BFP) tag, was expressed in Rosetta 2 cells and lysed in 100 mM Tris-HCl, pH 8.0, 450 mM NaCl, 1 mM MgCl₂, 1 mM EDTA, 0.5 mM PMSF, 6 mM BME, 200 μM GTP, 1 cOmplete EDTA-free protease inhibitor, and 1,000 U DNase I. Affinity purification was performed using a StrepTrap HP 5 ml column (GE Healthcare) with a binding buffer containing 100 mM Tris-HCl, pH 8.0, 450 mM NaCl, 1 mM MgCl₂, 1 mM EDTA, 0.5 mM PMSF, 6 mM BME, and 200 μM GTP. The protein was eluted with 100 mM Tris-HCl, pH 8.0, 450 mM NaCl, 1 mM MgCl₂, 1 mM EDTA, 0.5 mM PMSF, 6 mM BME, 200 μM GTP, and 2.5 mM d-desthiobiotin, then dialyzed overnight into CSF-XB supplemented with 10% (w/v) sucrose.

#### Tubulin labeling

Tubulin from bovine brain (PurSolutions) was labeled with Atto 647 NHS ester (GE Healthcare) following previously reported procedures^51^.

#### Kinesin-1 (K401) purification

K401, a truncated form of *Drosophila melanogaster* kinesin-1, was expressed in *Escherichia coli* (Rosetta 2 DE3) from plasmid DNA (AddGene, 15960). The K401 construct corresponds to the first 401 amino acids of kinesin-1, encompassing the motor domain and neck linker, and forms a truncated, dimeric motor capable of processive motility. A BCCP tag at the N-terminus enables site-specific biotinylation. Cells grew to an optical density of 0.6 and were induced with IPTG overnight. Following lysis, the clarified lysate was purified using nickel-affinity chromatography under conditions consistent with RanQ69L purification, and eluted fractions were analyzed to confirm protein integrity. The final concentration of purified K401 was 1.6 mg/ml, and its activity in MT gliding assays was ∼90%, with an average gliding speed of ∼330 nm/s.

#### Motor complex preparation

Quantum dot cargoes were prepared by incubating purified K401 with streptavidin-coated Qdot 525 (Thermo Fisher) at a final concentration of 0.1 µM for 1 hour at room temperature^66^.

### 4.2. Branched MT network gliding assay

Flow chambers were built by placing parallel strips of double-sided tape between a glass microscope slide and a cleaned glass coverslip to create a sealed channel with an approximate volume of 10 to 20 µL. To functionalize the surface, 1.5 mg/mL Neutravidin solution was introduced into the flow channel and incubated on ice for 20 minutes to allow adsorption to the glass surface. The channel was then washed thoroughly with BRB80 buffer supplemented with bovine serum albumin (BSA) to remove unbound Neutravidin and block nonspecific binding sites. BRB80 buffer consisted of 80 mM PIPES adjusted to pH 6.8 with KOH, 1 mM MgCl₂, and 1 mM EGTA. For surface blocking, BRB80 was supplemented with BSA at a final concentration of 5 mg/mL.

Biotinylated K401 motor protein was then introduced into the chamber and incubated at room temperature for 3 minutes to allow binding to the Neutravidin coated surface. The channel was subsequently washed with BRB80 containing 5 mg/mL BSA to remove unbound motors. A second aliquot of biotinylated K401 was flowed into the chamber and incubated for an additional 3 minutes at room temperature to increase motor surface density, followed by a gentle wash with blocking buffer.

Xenopus egg extract containing RanQ69L (10 µM), fluorescently labeled tubulin, and EB1-GFP (1 µM) was then introduced into the chamber. Branched MT network formation and gliding dynamics were imaged immediately after introducing the extract. For experiments in which transport of motor complexes was examined, 0.5 µL of motor complex solution was added to 10 µL of egg extract immediately prior to flowing the mixture into the chamber.

### 4.3. Microscopy and image processing

#### Ring TIRF microscopy

Imaging was performed on a Nikon Ti2 inverted microscope with a 100× oil immersion objective lens (1.49 NA) configured for ring TIRF microscopy. Fluorescence images were acquired using an ORCA Fusion BT sCMOS camera (Hamamatsu). Image acquisition and dual channel imaging were controlled using NIS Elements software (Nikon). For each experiment, exposure settings and acquisition parameters were optimized to maximize signal to noise while minimizing photobleaching. Images were typically acquired at a rate of 1 frame per second. Brightness and contrast were adjusted uniformly for visualization and analysis as appropriate.

#### Tracking

EB1 comets and motor complexes were tracked using the TrackMate plugin in Fiji (ImageJ). Detection settings were adjusted as needed for each experiment, and tracks were checked manually to remove incorrect detections. Translational speeds were calculated from the measured displacements using the calibrated pixel size and the imaging rate.

#### Optical flow analysis

Optical flow was computed in MATLAB using the Farnebäck method to compute dense velocity fields from time-lapse image sequences. Before calculating the flow, images were smoothed with a Gaussian filter with a threshold of 2 to reduce noise. The resulting flow vectors were converted into unit vectors to represent the local direction of motion independent of speed. To reduce fluctuations, the direction fields were coarse-grained by averaging the optical flow over every five consecutive frames.

## Acknowledgments

This work was supported by the Omenn–Darling Postdoctoral Fellowship, the Princeton University Eric and Wendy Schmidt Transformative Technology Fund, and NSF Grant No. DMS/NIGMS 2245850 (to H.A.S. and S.P.). The authors thank the members of the Petry Lab and Stone Lab for insightful discussions throughout this project.

## Author contributions

M.Z., N.S.W., H.A.S., and S.P. conceived the study and contributed to manuscript preparation. M.Z. developed the methodology, performed the investigation, and generated the visualizations. H.A.S. and S.P. acquired funding, administered the project, and supervised the research. All authors contributed to writing the manuscript.

## Competing interests

The authors declare no competing interests.

## Notes

### Competing Interest Statement

The authors have declared no competing interest.

